# Growth/differentiation factor 15 controls primary cilia morphology in the murine ventricular-subventricular zone thereby affecting progenitor proliferation

**DOI:** 10.1101/2023.08.30.555481

**Authors:** Katja Baur, Şeydanur Şan, Gabriele Hölzl-Wenig, Claudia Mandl, Andrea Hellwig, Francesca Ciccolini

**Author notes:** **Correspondence** should be addressed to: Dr. Francesca Ciccolini.

## Abstract

Growth/differentiation factor 15 (GDF15) and its receptor GDNF Family Receptor Alpha-Like (GFRAL) are expressed from embryonic development onwards in the germinal epithelium of the ganglionic eminence (GE), regulating proliferation and number of apical progenitors. However, the mechanisms underlying this regulation are not yet clear. We here show that GDF15 exerts this regulation by affecting ciliary signalling. Not only was GFRAL localized to primary cilia but, constitutive GDF15 ablation also led to shorter and thicker primary cilia. Lack of GDF15 affected the expression of histone deacetylase 6 (HDAC6) and ciliary adenylate cyclase 3 (ADCY3), thereby modifying acetylation of microtubules and endogenous Sonic Hedgehog (SHH) activation in neural progenitors. Application of exogenous GDF15 or pharmacological antagonism of HDAC6 or ADCY3 all increased cilia length and rescued proliferation and SHH signalling in mutant but not WT progenitors. Notably, HDAC6 expression and cilia length were changed only in the GE, were ciliary GFRAL localization was observed. In contrast, GFRAL was absent from primary cilia of hippocampal progenitors where GDF15 affected ADCY3 and SHH signalling, but not HDAC6 expression or cilia morphology. We conclude that ciliary GDF15 signalling regulates HDAC6 thereby affecting primary cilia elongation and proliferation in apical progenitors.

## Introduction

Growth and differentiation factor 15 (GDF15) is a divergent member of the transforming growth factor (TGF) β superfamily and it was recently found to regulate body weight by controlling food uptake via its receptor GDNF family receptor alpha-like (GFRAL). The latter it is scarcely expressed in the healthy adult brain where it is mainly observed by neurons in the brain stem and can induce signal transduction upon physical interaction with the receptor tyrosine kinase Ret (Emmerson et al. 2017; Yang et al. 2017). However, because it is ubiquitously expressed, GDF15 is associated not only with obesity, but also with a variety of diseases such as cancer and pathologies of the liver, kidneys and the cardiovascular system (Rochette et al. 2020). In the uninjured rat brain, high levels of GDF15 are observed around the lateral ventricle (Böttner et al. 1999; Schober et al. 2001), suggesting that the factor may affect adult neural stem cells (NSCs) in the ventricular-subventricular zone (V-SVZ), which along with the subgranular zone (SGZ) of the dentate gyrus in the hippocampus (HP) represent the main neurogenic niches in the adult mammalian brain. Consistent with an effect of GDF15 on NSC function, GDF15 is expressed in both neurogenic regions (Schober et al. 2001), where it promotes epidermal growth factor receptor (EGFR) expression and affects neural stem cells (NSC) proliferation (Carrillo-Garcia et al. 2014). However, unlike the diffused expression observed in the HP, in the V-SVZ GDF15 and its receptor are particularly expressed at the apical side contacting the lateral ventricle (Schober et al. 2001). Moreover, the V-SVZ differs from the HP also in the fact that in this region GDF15 prevents proliferation of apical progenitors from late development onwards. This effect is independent of changes in EGFR expression, observed in mutant progenitors, and it leads to increased number of apical cells in the adult V-SVZ, including ependymal cells and apical NSCs (Baur et al. 2023).

Notably, neural precursors contacting the lateral ventricle extend a primary cilium, which is important to sense and transduce environmental signals, such as sonic hedgehog (SHH) signalling, as well as to regulate cell cycle progression (Plotnikova, Pugacheva, and Golemis 2009; Khatri et al. 2014). Consistent with this, recent studies have highlighted that actively proliferating NSCs dismantle the organelle before undergoing cell division and the presence and the length of the organelle positively correlate with quiescence and cell cycle progression (Plotnikova, Pugacheva, and Golemis 2009; Khatri et al. 2014). In the adult V-SVZ, apical NSCs represent the most abundant population of ciliated progenitors, as the vast majority of NSCs present at the basal side of the neurogenic niche lack the organelle (Baur et al. 2022).

The activity of histone deacetylase 6 (HDAC6), which is responsible for deacetylation of ciliary microtubules, is of primary importance for dismantling and maintenance of primary cilia in concomitance with cell cycle progression (Sanchez de Diego et al. 2014; Shi et al. 2021). However, HDAC6 is not strictly a ciliary protein and although it is primarily localized in the cytoplasm its activity depends on its subcellular localization (Wang et al. 2016). In contrast to HDAC6, the adenylate cyclase 3 (ADCY3) is essentially localized to primary cilia and it has been shown to influence primary cilia length and SHH signalling by regulating cAMP levels (Bishop et al. 2007; Ou et al. 2009; Qiu et al. 2016). Interestingly, alteration in ciliary function like modulation of ADCY3 and GDF15, have all been associated with regulation of energy homeostasis, food intake and obesity (DeMars et al. 2023).

In light of the peculiar expression of GDF15 at the apical niche side and its effect on the proliferation of apical progenitors, we have here investigated its effect on primary cilia and the mechanism underlying this effect. Our data show that GDF15 critically contributes to primary cilia regulation in apical progenitors and NSCs in the embryonic GE and in the adult V-SVZ and that this effect correlates with ciliary localization of GFRAL and altered expression/function of HDAC6.

## Materials and Methods

### Animals

All animal experiments were approved by the Regierungspräsidium Karlsruhe and the local authorities at the University of Heidelberg. The Gdf15^−/−^ line was previously described (Strelau et al. 2009) and bred with homozygous animals with regular blood refreshing. For E18 matings, animals were mated for 48 hours and then separated, or separated after 24 hours when a vaginal plug was visible. The separation day was considered as E1.

### Whole mount dissection and incubation

Whole lateral walls were dissected and fixed as described before (Mirzadeh et al. 2008; Monaco et al. 2019). In short, the mice were killed by CO_2_-inhaltation and subsequent cervical dislocation (adult) or by decapitation (E18), and the brain was dissected in a buffered sucrose solution. The lateral wall was thinly removed and either directly fixed in ice-cold 3% paraformaldehyde (PFA), 4% sucrose in phosphate-buffered saline (PBS), or incubated at 37°C overnight in 1 ml Euromed-N (Euroclone, #ECM0883L) containing 2% B27 (Invitrogen, #17504044) and, when indicated, recombinant human GDF15 (10 ng/ml; R&D Systems, Q99988), human recombinant EGF (20 ng/ml; Peprotech, #AF-100-15), PD158780 (20 µM; Calbiochem/Merck, #513035) or AMD3100 octahydrochloride hydrate (6 µM; Sigma Aldrich, #A5602), Tubastatin A (10 µM; Selleckchem, #S8049), NKY80 (200 µM; Sigma Aldrich, #116850), U0126 (10 µM; Enzo Life Sciences, #BML-EI282-0001), Smoothened agonist (SAG; 200 nM; Cayman Chemicals, #11914). The incubated lateral walls were fixed the next day, and all dissections were left in the 3% PFA, 4% sucrose in PBS solution at 4°C overnight, then kept in PBS containing 0.01% azide until immunostaining.

### Brain sections

For brain sections, mice were killed as for the whole mount; subsequently, the entire brain was removed and fixed in 4% PFA in PBS for 72 hours at 4°C with slight agitation. For cryoprotection, the brains were afterwards submerged in a 30% sucrose in PBS solution for 48 hours. The cryoprotected brains were then frozen at −20°C and sliced in 20 µm thick sections using a Leica CM1950 Cryostat. Coronal or sagittal slices were rinsed in PBS and stored in PBS containing 0.01% azide until immunostaining.

### Immunostaining

Immunostaining of brain sections and whole mounts was performed as previously described (Luque-Molina et al. 2019; Monaco et al. 2019). For IdU staining, the whole mounts were incubated in 2N HCl for 30 min at 37°C, followed by 30 min of 0.1 M Na-tetraborate pH 8.5 at RT, before the standard protocol. For antibodies used, please refer to supplementary table S1.

### Scanning Electron Microscopy (ScEM)

Scanning electron microscopy was performed as described before (Monaco et al. 2019).

### Real-time qPCR

For real-time quantitative PCR (qPCR), RNA was extracted using a Quick-RNA Microprep kit (Zymo Research). For this, tissue (SVZ or HP from one hemisphere, or brain stem and hypothalamus, dissected as described above) was lysed in 300 µl lysis buffer; the RNA was extracted according to manufacturer’s instructions, including a DNase digestion step. The RNA was eluted from the column in 20 µl RNase-free water, and then retrotranscribed into cDNA using M-MLV reverse transcriptase (Promega) and Oligo(dT)15 primers (Promega) according to manufacturer’s instructions in a ProFlex PCR machine (Applied Biosystems).

1 µl cDNA was used for qPCR in a volume of 20 µl using TaqMan GeneExpression Master Mix (Thermo Fisher Scientific) according to manufacturer’s instructions using a StepOne Plus Real-Time PCR System (Thermo Fisher Scientific). For probes used, please refer to supplementary table S2.

For normalization, beta-actin was used as a housekeeping gene; fold change was calculated as 2^-ΔΔCT^, where ΔC_T_ is the difference between cycle threshold (C_T_) value of the gene of interest to its respective beta-actin C_T_ value, and ΔΔC_T_ is the difference between the ΔC_T_ value of a control sample (e.g. wild-type untreated) and that of the sample of interest. All samples were run as duplicates.

### Flow cytometry

Flow cytometry was performed as previously described (Luque-Molina et al. 2019; Baur et al. 2022). Cells were labelled with rat-anti-Prominin-1 antibody (Brilliant Violet 421-conjugated; BioLegend, #141213, Lot# B255870, 1:1000) and Alexa Fluor 647-conjugated recombinant human EGF (Invitrogen, #35351, Lot# 1700382, 1:1000) to label EGFR.

### Imaging and Data analysis

The immunolabelled tissues were imaged using a Leica SP8 confocal microscope and analysed with Fiji/ImageJ (Schindelin et al. 2012); calculations were done using Microsoft Excel. To measure the cilium length, cilia were marked on projections of confocal z-stacks using the ImageJ straight line or segmented line selection tool; the line was drawn from end to end along the centre of the ciliary axoneme. For thickness, lines were drawn perpendicular to the cilium at the point where the individual cilium was the thickest. To assess the number of cilia, cilia in the V-SVZ were counted only if they were above the apical surface, i.e. without visible DAPI staining, inside one representative 50 µm x 50 µm square per image and normalized on this area. In the DG, HTh and BSt, cilia in all layers of a z-stack were counted in a projection image and normalized on the number of nuclei. All cilia, and the nuclei in the DG were counted manually using the “Cell Counter” ImageJ plugin, while nuclei in the HTh and BSt were counted using an ImageJ-macro utilizing the “Analyse Particles” function as well as the “Adjustable Watershed” plugin.

Apical and subapical dividing cells were scored as described before (Baur et al. 2023).

### Statistical analysis and graphs

Statistical analysis and graphing were performed with GraphPad Prism 8. Statistical significance was determined by a two-tailed Student’s t-test for two, and by two-way ANOVA with Dunnett’s multiple comparisons test for more than two groups. Significance was reached at *p<0.05, **p<0.01, ***p<0.001. All bars represent the mean value ± standard error of the mean (SEM). The data points indicate values of individual animals, i.e. the n number is represented by the respective number of data points in each bar.

## Results

### GDF15 ablation changes cilia morphology in the embryonic GE and adult V-SVZ

In a previous study, we found that both GDF15 and its receptor GFRAL are expressed at the apical side of the adult V-SVZ and in the corresponding germinal niche of the ganglionic eminence at embryonic day 18 (E18) (Baur et al. 2023). Since apical cells are characterized by the presence of primary cilia, we investigated the possibility that GFRAL is also present in the organelle. Closer examination of the cilia in E18 wild-type (WT) embryos confirmed the presence of the receptor in cilia protrusions identified by immunofluorescence with antibodies against the ADP-ribosylation-like factor 13b (Arl13b) and adenylate cyclase 3 (ADCY3, also known as AC3), both expressed in primary cilia at this age (Monaco et al. 2019). We found that in the apical GE, some Arl13b- and especially ADCY3-expressing cilia were enriched in GFRAL (Fig 1A, white arrows). We also detected a few cilia with only Arl13b and GFRAL (blue arrows), or ADCY3 and Arl13b without GFRAL (turquoise arrows). Notably, the receptor was also observed in primary cilia of the adult V-SVZ identified by Alr13b staining (supplementary Fig. S1A). In contrast, a similar analysis on coronal sections of the E18 HP revealed only a rather weak expression of GFRAL and no co-localization with Arl13b or ADCY3 (Fig 1A), both of which are widely expressed by primary cilia in this region at E18 (Berbari et al. 2007; Caspary, Larkins, and Anderson 2007).

**Figure 1:**
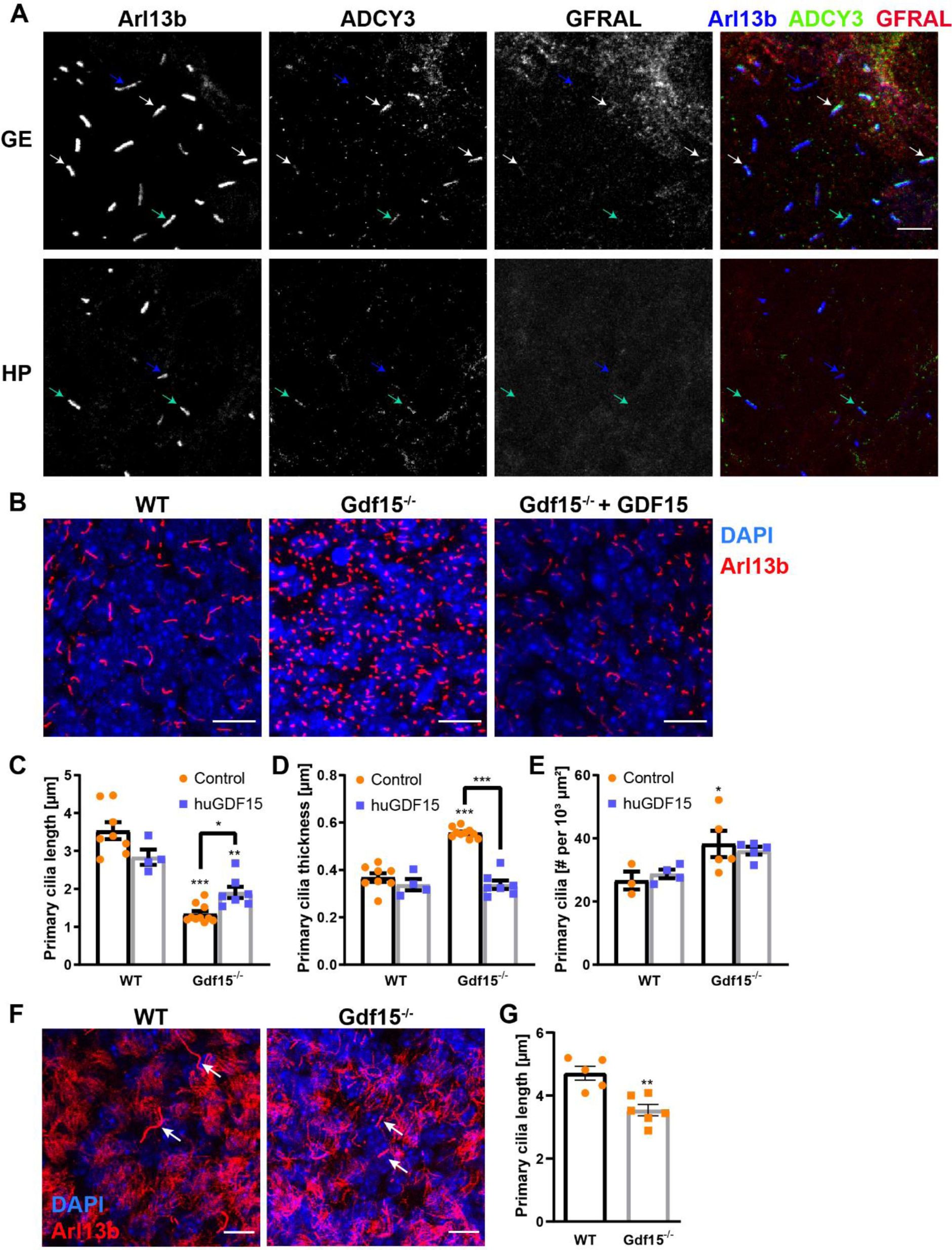
GDF15 affects cilia morphology and number in the lateral periventricular germinal zone. (A) Representative confocal micrographs of the apical ganglionic eminence (GE) and of the prospective dentate gyrus in the hippocampus (HP) obtained from whole mount preparations of E18 embryonic brain tissue upon triple immunostaining with ciliary marker Arl13b (blue), ADCY3 (green) and GFRAL (red). White arrows point at cilia expressing both ciliary markers and GFRAL, green and blue arrows to cilia expressing both ciliary markers, or Arl13b only, respectively. Scale bar = 5 µm. (B, F) Representative confocal micrographs of whole mount preparations of the GE obtained from E18 embryos (B) and adult V-SVZ (F) of the given genotype and treated with 10 ng/ml exogenous GDF15 as indicated. Tissue preparations were immunostained with antibodies to aArl13b (red) and DAPI (blue) counterstaining of the nuclei to visualize apical cilia. Scale bars = 10 µm. (C-E, G) Quantification between genotypes and treatment in length (C, G), thickness (D) and number (E) of primary cilia of E18 (C-E) and adult (G) animals. Bars indicate mean ± SEM. Each data point represents summarized data from one individual animal. *indicates significance (*p<0.05, **p<0.01, ***p<0.001) compared to WT (on top of bars) or between treated and untreated sample (on top of line).

Proteins expressed in primary cilia often have a role not only in ciliary signalling, but they are also vital for ciliary morphology and organelle function. Ablation of such proteins often leads to less, malformed, or dysfunctional cilia (Bachor et al. 2017; Chen et al. 2016; Ehnert et al. 2017; Escudier et al. 2009; Frasca et al. 2020; Larkins et al. 2011; Lechtreck et al. 2008; Nakakura et al. 2015; Ou et al. 2009; Patnaik et al. 2019; Pazour et al. 2002). Therefore, we compared the structure of primary cilia in the V-SVZ of WT and Gdf15-knock-out/LacZ-knock-in (Gdf15^−/−^) E18 embryos using immunofluorescence for the ciliary protein Arl13b (Fig. 1C). This analysis revealed that primary cilia in Gdf15^−/−^ embryos are on average shorter (Fig. 1D), thicker (Fig. 1E) and more numerous (Fig. 1F) than in WT age-matched controls. Whereas cilia in the WT V-SVZ were on average around 3.67 ± 0.45 µm long, 0.39 ± 0.02 µm thick and about 14.96 ± 1.98 cilia per 1000 µm², in Gdf15^−/−^ embryos, they were only 1.84 ± 0.12 µm long, 0.54 ± 0.01 µm thick, and 23.44 ± 3.89 cilia per 1000 µm² were detected. To exclude that these structural defects are artefacts of immunofluorescence due to differential protein expression, we investigated cilia morphology using scanning electron microscopy (ScEM) (Supplementary Fig. S1B). Overall cilia length measured in ScEM micrographs was consistent with the values measured in immunofluorescently labelled samples (Supplementary Fig. S1B) whereas the overall thickness was decreased (0.19 µm ScEM compared to 0.39 µm in fluorescence). However, both average values were significantly smaller in Gdf15^−/−^ compared to WT animals.

Underscoring the direct role played by the growth factor in regulating cilia morphology, incubation of the dissected embryonic GE with exogenous human GDF15 for 24 hours led to an amelioration of the ciliary defect in the mutant V-SVZ in terms of length and thickness and to a normalization of the number of primary cilia to WT levels (Fig. 1D-F).

While GDF15 is not highly expressed in early embryonic stages until E14, it is upregulated around E16/E18 and it remains at a similar level in the postnatal brain (Carrillo-Garcia et al. 2014). Therefore, we hypothesized that the effect of the lack of GDF15 might persist into adulthood, so we next examined primary cilia length at the apical side of the V-SVZ of adult WT and Gdf15^−/−^ mice (Fig. 1G). Whereas primary cilia in adult progenitors of Gdf15^−/−^ animals measuring 3.54 ± 0.18 µm were longer than the embryonic counterpart, they were still significantly shorter than primary cilia of progenitors in WT mice, which were on average 4.71± 0.22 µm long (Fig. 1H, white arrows).

To investigate whether GDF15 affects cilia morphology in other brain regions we measured cilia length and number in the HP, hypothalamus and brain stem. These brain regions were selected because albeit in all of them a function for GDF15 and the presence of GFRAL was demonstrated (Yang et al. 2017; Schober et al. 2001), in these regions GFRAL did not display ciliary localization. Using Arl13b and ADCY3 to visualize cilia in the E18 (Supplementary Fig. S2A, B) and adult HP (Supplementary Fig. S2C-E), as well as adult hypothalamus (Supplementary Fig. S2F-H) and brain stem (Supplementary Fig. S2I-K), both Arl13b^+^ or ADCY3^+^ cilia we found no change in either length (Supplementary Fig. S2B, D, G, J) or number of cilia per cell (Supplementary Fig. S2E, H, K) in these brain regions between Gdf15^−/−^ and WT controls animals. The number of cilia per cell were not counted in the E18 hippocampus, as the nuclei were too dense to count at this age. These data suggest that the effect of GDF15 on the primary cilia might due to ciliary GFRAL.

Upon exposure to its ligand, GFRAL dimerizes and binds to co-receptor RET, which activates several signalling cascades, among them the MAPK-ERK pathway (Mullican et al. 2017). Therefore, to further support the hypothesis that the effect of GDF15 on cilia depends on GFRAL activation, we investigated its dependence from ERK activation and transcription. To determine whether the morphological rescue upon GDF15 application requires ERK activation, we used MEK1/2-inhibitor U0126 to inhibit phosphorylation of ERK. Analysis by western blot of whole E18 GEs incubated in medium without or with either GDF15 or epidermal growth factor (EGF), which also activates the MAP-ERK pathway, for 24 h in the presence or absence of additionally U0126 to inhibit ERK phosphorylation. As expected, both GDF15 and EGF induced ERK phosphorylation, which was diminished when U0126 was present. Parallel samples were not analysed by western blot, but instead fixed and processed for cilia analysis by immunofluorescence (Fig. 2B, D). Here we found that GDF15 application did not lead to cilia lengthening while MEK1/2 was inhibited, consistent with the fact that the effect is mediated by GFRAL via ERK signalling. Moreover, activation of ERK signalling by EGF was not sufficient to rescue cilia morphology, whereas blockade of transcription by actinomycin D did not affect cilia morphology nor the ability of GDF15 to rescue the morphological defects in mutant primary cilia (Fig. 2B, D). Taken together, these data show that GDF15-GFRAL dependent activation of MAPK/ERK signalling, but not transcription, is necessary for the effect of GDF15 on primary cilia.

**Figure 2:**
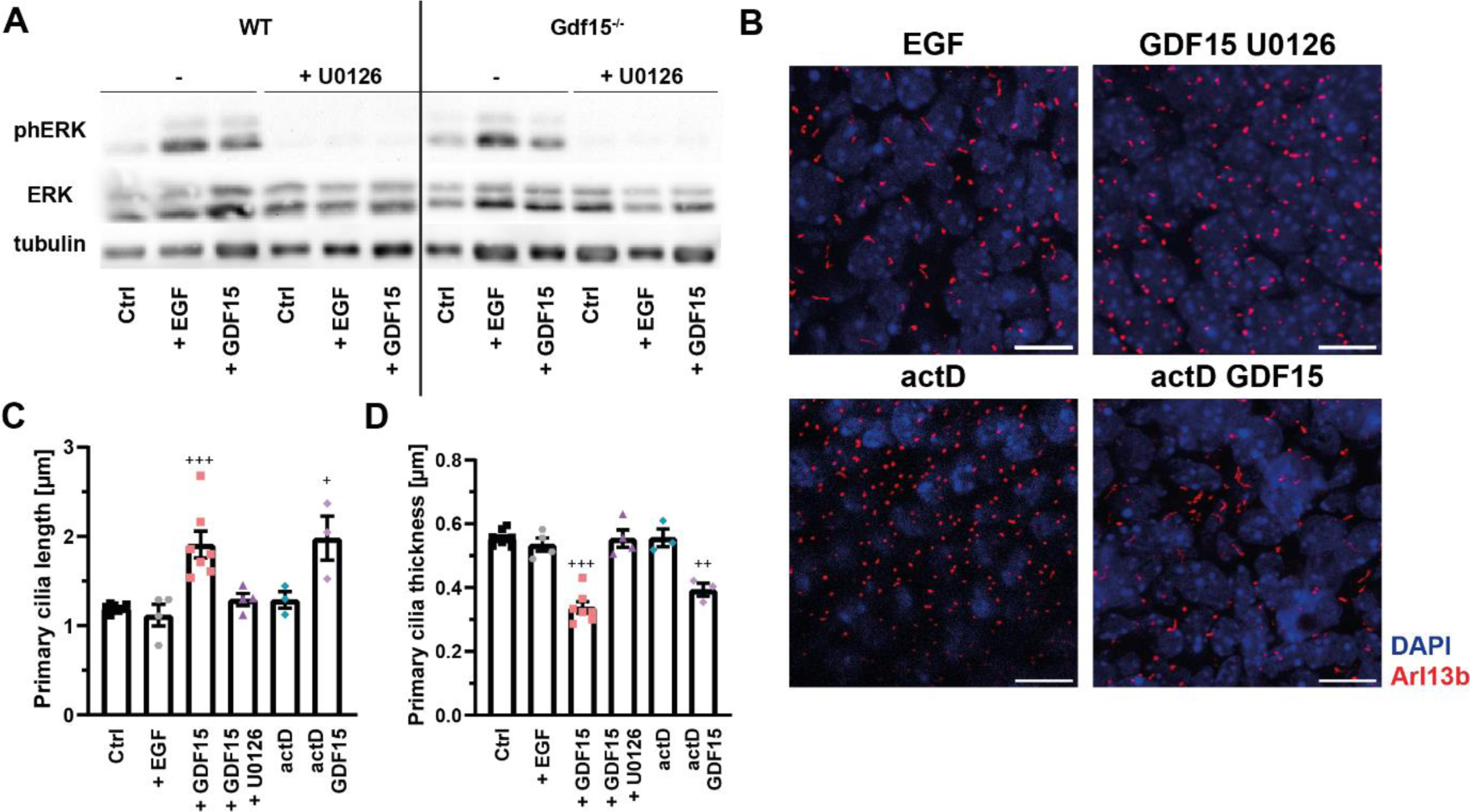
GDF15 effect on primary cilia requires ERK activation and is transcription independent. (A) Representative images of western blot analyses of total (ERK) and phosphorylated (phERK) ERK levels in the whole E18 ganglionic eminence (GE) of the indicated genotype cultured in the presence and absence of the given treatments for 24 hours. (B) Representative confocal micrographs of primary cilia at the apical side of whole mount preparations of the Gdf15^−/−^ E18 GE upon exposure to the given treatments for 24 hours. Thereafter the tissue was processed for immunostaining with Arl13b antibodies (red) and DAPI (blue) nuclear counterstaining. Scale bars = 10 µm (C, D) Quantification of the effect of the treatments on length (C) and thickness (D) of primary cilia. Bars indicate mean ± SEM. Each data point represents summarized data from one individual animal. ^+^ indicates significance (^+^p<0.05, ^++^p<0.01, ^+++^p<0.001) compared to untreated controls (Ctrl).

Cilia are known to relay important signals such as SHH signalling. To determine whether the changes in ciliary morphology impact ciliary signalling in the mutant mice, we analysed the levels of mRNA SHH effector gene *Gli1* in the V-SVZ of E18 WT and Gdf15^−/−^ animals using qPCR. We found that in control conditions (DMSO-treated), *Gli1* mRNA was reduced by half in the GE (Fig. 3A, left panel). However, a similar change was also observed in the HP (Fig. 3A, right panel) acutely dissected from Gdf15^−/−^ E18 embryos and treatment of the WT and mutant GE with smoothened agonist (SAG), a SHH pathway activator, led to a similar significant increase in *Gli1* transcripts in both tissues, indicating that changes in endogenous SHH activation are not dependent on cilia morphology, and that despite bearing shorter cilia, progenitors in the GE are still responsive to SMO relocation.

**Figure 3:**
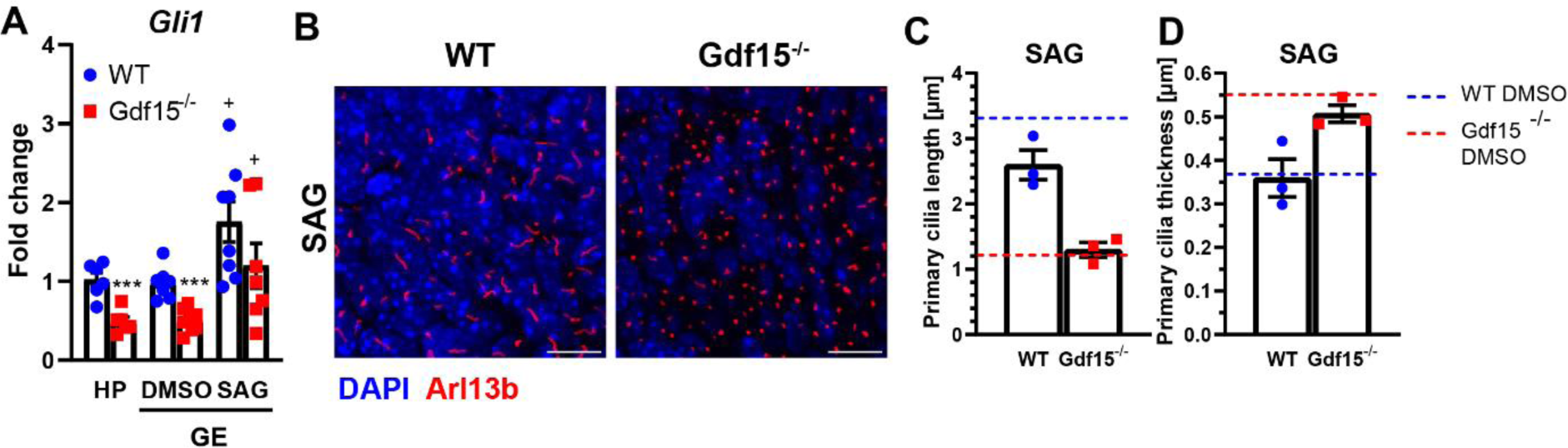
GDF15 affects endogenous SHH activation. (A) Fold-change in the expression of mRNA *Gli1* levels in the whole E18 hippocampus (HP) or ganglionic eminence (GE) of WT and Gdf15^−/−^ animals, the latter treated with DMSO as control or with smoothened agonist (SAG) as indicated. * and + indicate significance between the treatments and genotype respectively. (***p<0.001; ^+^p<0.05). (B) Representative confocal micrographs of primary cilia at the apical side of whole mount preparations of the E18 ganglionic eminence (GE) of the given genotype treated with SAG for 24 hours and processed for immunostaining with Arl13b antibodies (red) to visualize primary cilia and DAPI (blue) nuclear counterstaining. Scale bars = 10 µm. (C, D) Quantification of the effect of SAG on length (C) and thickness (D) of primary cilia, Values of DMSO-treated controls are illustrated by the blue (WT) and red (Gdf15^−/−^) dashed bars. Bars indicate mean ± SEM. Each data point represents summarized data from one individual animal.

Furthermore, after application of SAG, neither the length (Fig. 3C) nor the thickness of primary cilia (Fig. 3D) were changed compared to a DMSO-treated control. Thus, lack of GDF15 affects cilia morphology by a mechanism that is independent of SHH signalling.

### ADCY3 and HDAC6 like GDF15 control ciliary length

Besides endogenous SHH activation, EGFR surface expression and cell cycle progression are also affected in the apical GE of Gdf15^−/−^ E18 embryos (Baur et al. 2023). Therefore, we investigated whether the expression of ADCY3 and HDAC6 is altered in mutant progenitors. The first localizes predominantly in primary cilia, including in embryonic neural progenitors (Monaco et al. 2019), where it promotes the synthesis of cAMP, in response to activation of G-protein coupled receptors, thereby regulating cilia length (Ou et al. 2009) and SHH signalling (Ou et al. 2009). The second instead regulates α-tubulin acetylation and it is important for the shedding of primary cilia before mitosis but also for the intracellular trafficking of EGFR (Pugacheva et al. 2007; Sanchez de Diego et al. 2014). Quantitative analysis of mRNA levels of *Adcy3* and *Hdac6* by real-time PCR (qPCR) in the embryonic GE (Fig. 4A) and in the HP (Fig. 4B), revealed that compared to the WT counterpart tissue, transcripts for *Adcy3* were significantly increased in the absence of GDF15 in the HP and especially in the GE, whereas lack of GDF15 led to an increase in *Hdac6* expression in the GE but not HP. Moreover, exposing either tissues to exogenous GDF15 led to decrease in both messengers in the mutant E18 GE eminence but not in the WT counterpart or in the HP obtained by either genotypic group. In immunofluorescently labelled E18 whole mount preparations of the GE, we also found a visible increase in ADCY3 immunoreactivity in Gdf15^−/−^ animals compared to WT controls. This increase, however, was not observed in primary cilia of the HP (see supplementary Fig. S2) and in the GE was no longer detected upon application of exogenous GDF15 (Fig. 4C), indicating that the increase in ADCY3 immunoreactivity in the GE may reflect the shortening of primary cilia. These conclusions could not be further tested by western blot analysis due to the poor performance of the available ADCY3 antibodies. Thus, independent of GFRAL localization, lack of GDF15 leads to an increase in *Adcy3* transcripts, whereas change in *Hdac6* mRNA levels in the mutant tissue is associated with ciliary localization of GFRAL. To determine whether the overexpression of ADCY3 and HDAC6 underlies the change in cilia morphology in Gdf15^−/−^ animals, we used the pharmacological agents NKY80 and Tubastatin A (TBA) to inhibit function of ADCY3 and HDAC6, respectively. For this, E18 whole GE preparations were incubated with or without the agents for 24 hours and cilia length and thickness were assessed in immunofluorescent images using Arl13b as a ciliary marker (Fig. 4D). Here, we found that both inhibitors led to an increase in length and to a decrease in thickness in the cilia of the apical Gdf15^−/−^ GE, similar to the effect of GDF15 (Fig. 4E, F). However, unlike exposure to exogenous GDF15, the treatments also led to a change in cilia morphology in WT progenitors. Whereas NKY80 caused increased ciliary length also in WT progenitors, TBA led unexpectedly to a shortening of primary cilia in this cell group (Fig. 4E). Thus, whereas ADCY3 promotes a decrease in cilia length in both WT and mutant progenitors, the effect of HDAC6 activity on cilia length depends on the genotype. Since HDAC6 activity is associated to its local intracellular regulation (Li, Shin, and Kwon 2013), the opposite effect of TBA on the length of cilia in WT and mutant progenitors indicates that HDAC6 activity/location may vary between the two groups of progenitors.

**Figure 4:**
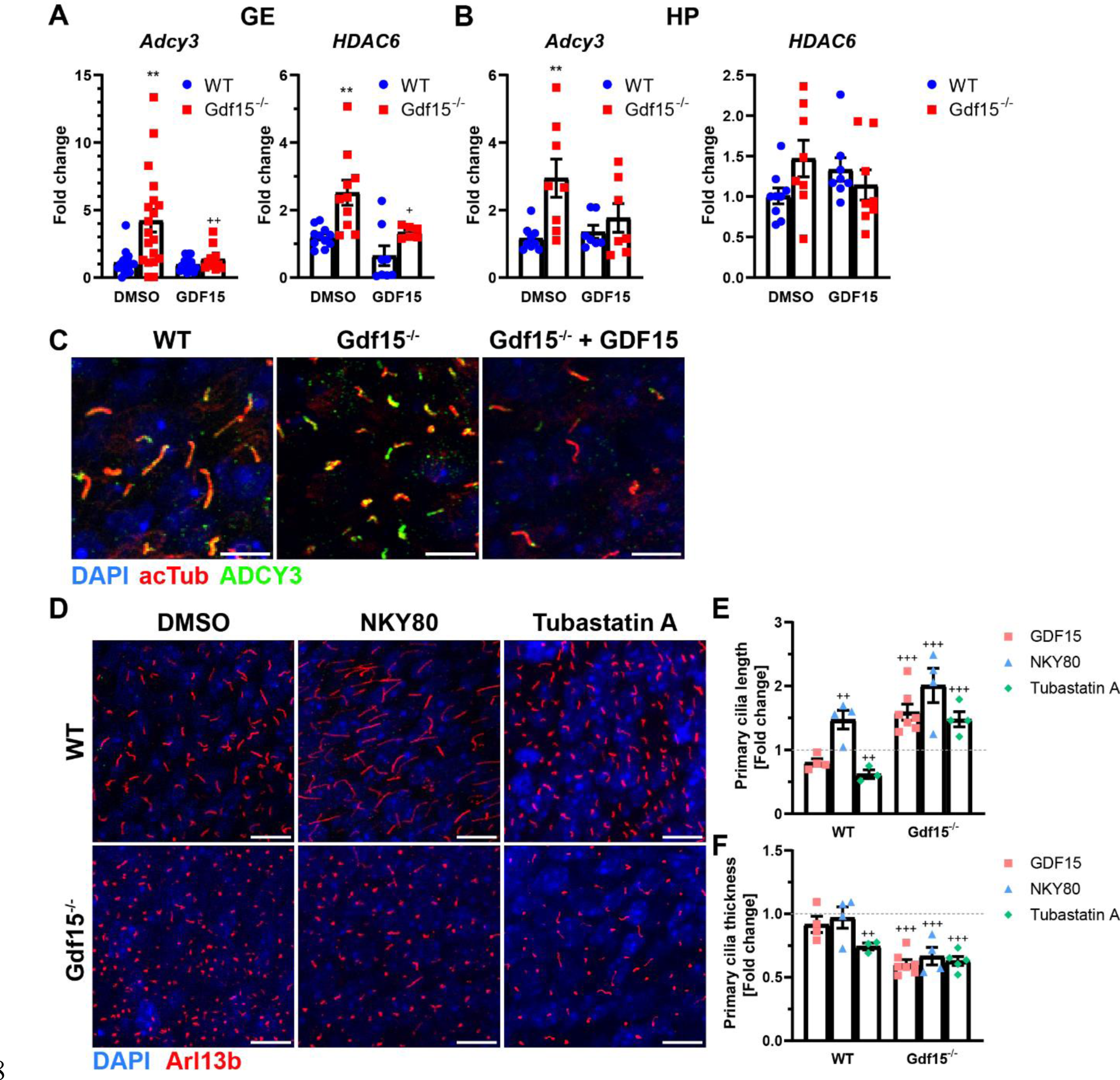
GDF15 modulates *Adcy3* and *Hdac6* expression. (A, B) Fold change in the expression of the given mRNAs in the whole GE (A) or HP (B) of E18 WT and Gdf15^−/−^ animals, normalized to the WT control. (C) Representative confocal micrographs of primary cilia at the apical side of whole mount preparations of the E18 ganglionic eminence (GE) of the given genotype upon immunostaining with as indicated. Note that the expression of ADCY3 but not acetylated tubulin (acTub) appeared to be modified according genotype/treatment. (D) Representative confocal micrographs of primary cilia at the apical side of whole mount preparations of the E18 GE of the given genotype treated as indicated for 24 hours and processed for immunostaining with Arl13b (red) antibodies to visualize primary cilia. DAPI nuclear counterstaining is visualized in blue. Scale bars indicate 10 µm. (E, F) Quantification of the effect of the treatment on cilia length (E) and thickness (F). Bars indicate mean ± SEM normalized to values measured in the WT untreated GE illustrated by dashed bars. Each data point represents summarized data from one individual animal. */^+^ indicate significance between Gdf15^−/−^ and WT animals (*) or between treated samples and DMSO-treated control (^+^) */^+^p<0.05, **/^++^p<0.01, ^+++^p<0.001.

Although exposure to exogenous GDF15 does not require transcription to rescue cilia morphology in mutant progenitors, we have observed that it modulates the expression of *Adcy3* and *Hdac6* transcripts in the mutant GE, showing a direct involvement of the growth factor in the regulation of the two enzymes. We therefore next investigated whether blockade of either enzyme reproduced a similar effect on transcript expression (supplementary Fig. S3B). This analysis revealed that in mutant tissue, both blockers had a similar effect on transcript levels as exposure to exogenous GDF15. However, both treatments reduced *Adcy3* but not *Hdac6* levels in WT progenitors (supplementary Fig. S3B), indicating that endogenous GDF15 signalling affects *Hdac6* expression upstream of either signalling molecule.

### Gdf15^−/−^ and WT apical progenitors display distinct tubulin acetylation patterns

HDAC6 regulates cilia length by deacetylation of ciliary tubulin, thereby shortening the ciliary axoneme, which is also known to affect cell cycle dynamics (Ehnert et al. 2017; Pugacheva et al. 2007; Shi et al. 2021; Sanchez de Diego et al. 2014). Interestingly, inhibition of HDAC6 caused lengthening of cilia only in mutant progenitors but not in the WT counterpart. Since HDAC6 is known to regulate cilia length also by modulating intracellular trafficking and transport, the differential effects observed in the two groups of progenitors may reflect the fact that HDAC6 is active in deacetylating microtubules at the ciliary and intracellularly in mutant and WT progenitors, respectively. To investigate this hypothesis, we used western blot to determine the overall levels of acetylated tubulin (acTub) in whole lateral walls of E18 WT and Gdf15^−/−^ animals, with and without incubation with HDAC6-inhibitor TBA, as well as exogenous GDF15 for 24 h (Fig. 5A, B). Here, we found that even though HDAC6 is overexpressed in Gdf15^−/−^ animals, there is no overall change to acTub protein levels between WT and mutant tissue even upon application of exogenous GDF15. In contrast, inhibition of HDAC6 by TBA, as expected, lead to a significant increase of acTub in the WT GE, but strikingly not in the Gdf15^−/−^ counterpart, where we detected only a non-significant trend increase in acTub levels (Fig. 5A, B). This lack of increase in overall tubulin acetylation in Gdf15^−/−^ animals, even though ciliary effects are clearly visible, might be due to differential localisation or activation of HDAC6. Therefore, we assessed levels of, HDAC6 protein in whole mount preparations using immunofluorescence (Fig. 5C, E). Here we found that treatment with TBA quantitively did not change the protein level as measured by fluorescence intensity, and there was no significant difference between WT and Gdf15^−/−^ samples (Fig. 5E). However, the punctuate staining visible in untreated samples was replaced by a more evenly distributed staining, suggesting a change in protein distribution upon HDAC6 inhibition. On the other side, analysis of acTub in these preparations confirmed the western blot analysis, clearly showing that TBA treatment increases acTub at the apical surface of the WT GE but not in the mutant counterpart (Fig. 5C, D). Taken together, these data indicate that the change in HDAC6 expression between WT and GDF15 may concern activation rather than localization of HDAC6.

**Figure 5:**
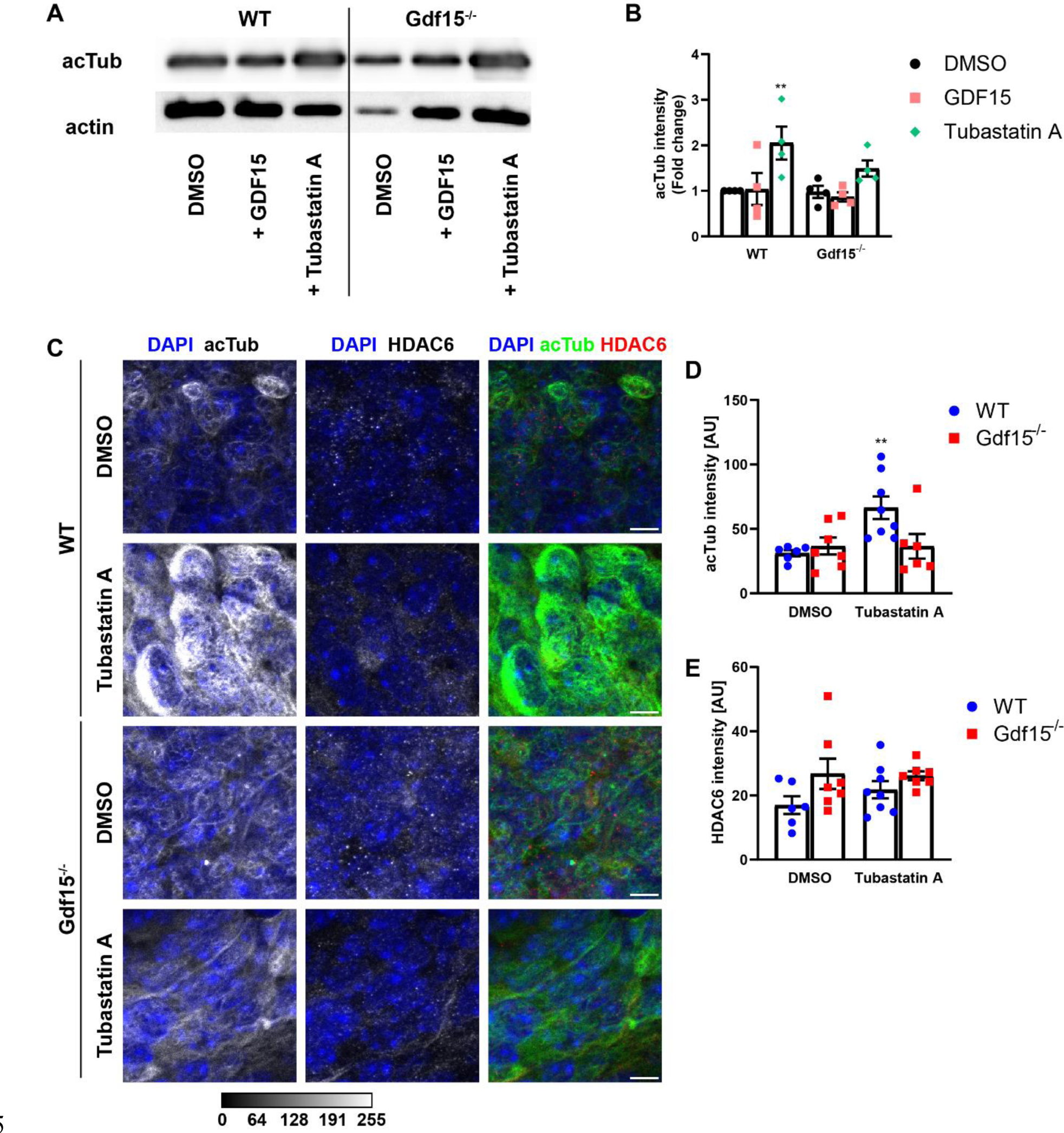
GDF15 modulates HDAC6 activity. (A) Western blot for acetylated tubulin (acTub) in the whole E18 ganglionic eminence (GE) of the indicated genotype incubated in the presence of the given treatments for 24 hours. (B) Quantification of acTub intensity in (A), normalized to actin. (C) Representative confocal micrographs of the apical side of whole mount preparations of the E18 GE of the given genotype and upon exposure to the indicated treatments for 24 hours. Thereafter the tissue was processed for immunostaining with antibodies to acTub (green), HDAC6 (red) and DAPI (blue) nuclear counterstaining. Scale bars = 10 µm. (D, E) Quantification of the effect HDAC6 blockade on acTub (D) and HDAC6 (E) fluorescence levels. Bars indicate mean ± SEM. Each data point represents summarized data from one individual animal. * indicates significance (**p<0.01) compared to DMSO-treated WT control.

### Effect of ciliary length and SHH on proliferation in the mutant apical GE

In a previous publication, we demonstrated that ablation of GDF15 leads to an increased proliferation of embryonic apical and subapical progenitors by increasing the number of cycling cells and by promoting cell cycle progression (Baur et al. 2023). A causal relationship between the defect in cilia morphology and proliferation is likely because the length of primary cilia regulates cell cycle progression (Malicki and Johnson 2017) and application of exogenous GDF15, similar to the altered ciliary morphology investigated here, was able to reverse the changes in proliferation. To test the causal relationship between ciliary and proliferation phenotype in mutant progenitors, we used the pharmacological blockers for HDAC6 and ADCY3 to determine if blockade of their activity leads to a decrease in cell cycle speed. For this, we used immunofluorescence for cell cycle marker Ki67 to label cycling cells (Fig. 6A) and looked at nuclear morphology or Ki67^+^ nuclei to determine which cells were currently undergoing mitosis (mitotic cells; Fig. 4A, white arrow). Using this approach, we found that in Gdf15^−/−^ animals, both TBA and NKY80 led to a decrease in the number of total cycling (Fig. 4B) and mitotic cells (Fig. 4C) in the E18 GE, similar to the effect of GDF15. Thus, blockade of HDAC6 and ADCY3 in mutant progenitors recapitulate the effect of GDF15 on cilia morphology and proliferation, indicating that GDF15 ablation leads to altered proliferation by affecting ciliogenesis. In contrast, in the WT tissue either treatment did not lead to a change in proliferation, either cell cycle or mitosis. This shows that activity of ADCY3 and HDAC6, as well as changes in ciliary length per se, are not enough to modulate proliferation of neural progenitors.

**Figure 6:**
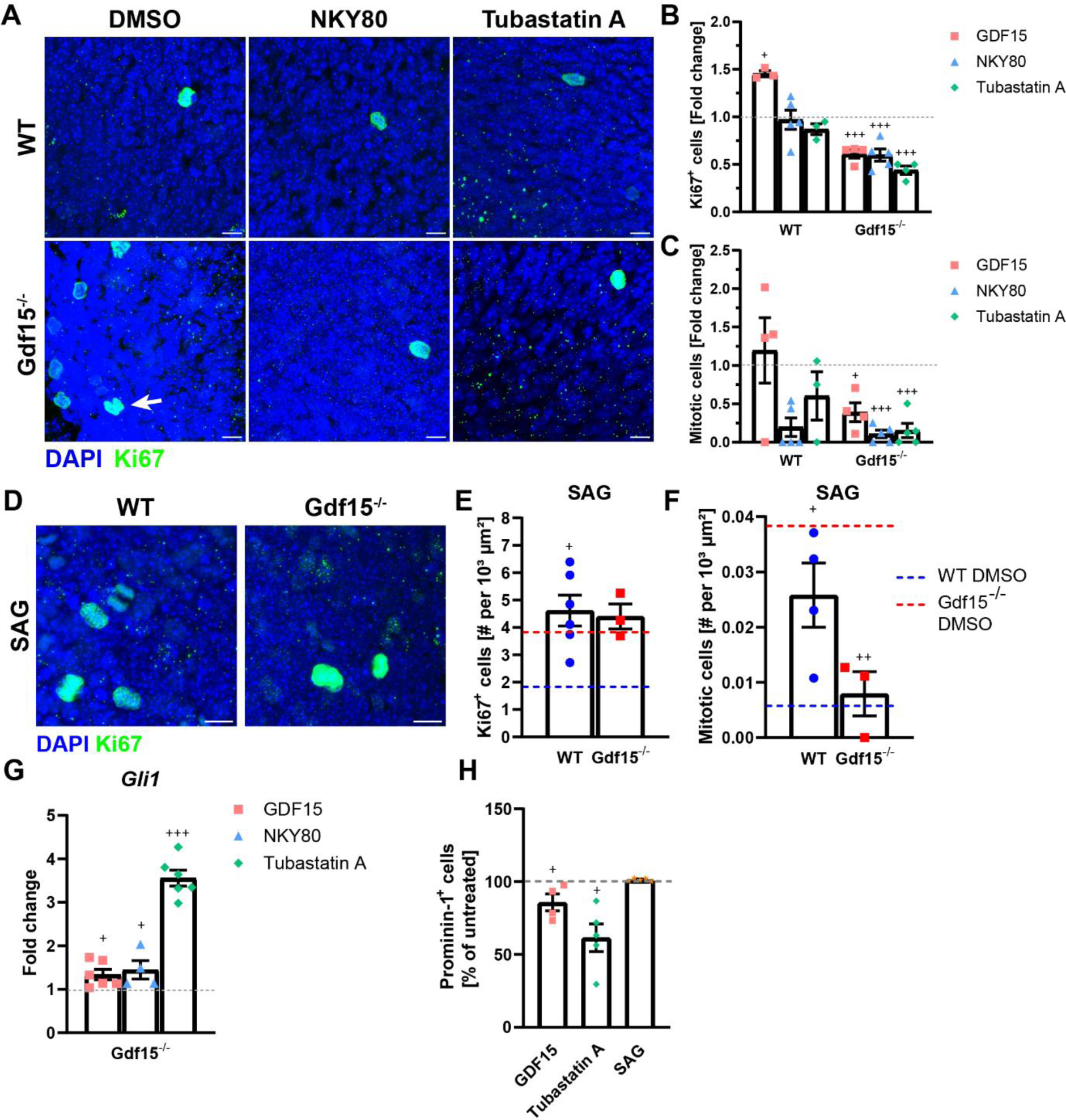
SHH modulates cell cycle length. (A,D) Representative confocal micrographs of cycling cells at the apical side of whole mount preparations of the E18 ganglionic eminence (GE) of the given genotype. Tissue was exposed to NKY80 or Tubastatin A (A) or SAG (D) for 24 hours and processed for immunostaining with Ki67 (green) antibodies and DAPI (blue) nuclear counterstaining. Scale bars indicate 10 µm. (B,C) Fold change of total cycling (B) and apically dividing (C) cells, normalized to respective DMSO-treated controls (dashed line). (E, F) Quantification of total cycling (E) and apically dividing (C) cells. Values of DMSO-treated controls are illustrated by the blue (WT) and red (Gdf15^−/−^) dashed lines. (G) Fold change in the expression of *Gli1* mRNA in the E18 mutant GE after treatment with indicated agents. Values are normalized to the respective untreated sample (dashed line). (H) Quantitative analysis of the number of Prominin-labelled cells in the Gdf15^−/−^ GE treated as indicated, as determined by flow cytometry. Values represent the percentage of the number of Prominin-1 immunopositive cells compared to DMSO-treated controls (dashed line). Bars indicate mean ± SEM. Each data point represents summarized data from one individual animal. + indicates significance from respective DMSO-treated control; ^+^p<0.05, ^++^p<0.01, ^+++^p<0.001.

Since mutant progenitors display impaired SHH signalling, which may also affect proliferation, we next investigated the effect of SAG on progenitor proliferation. Consistent with previous observations (Komada 2012) activation of SHH signalling in WT mice increased the number of both cycling and dividing cells (Fig. 6D-F). In Gdf15^−/−^ embryos, SAG instead led to a significant decrease only of mitotic cells (Fig. 6F) without affecting the number of total cycling cells (Fig. 6E), indicating that impaired endogenous SHH activation in mutant progenitors is responsible for the increase in the number of cells undergoing mitosis but not for the increase in cell cycle re-entry.

We next investigated whether inhibition of HDAC6 and ADCY3, like exposure to exogenous GDF15, also affect *Gli1* expression (Fig. 6G). Since we have seen that both blockers affect proliferation only in the mutant GE, we performed these experiments only in this group of progenitors. We found that that NKY80 and TBA all led to a significant increase in *Gli1* transcripts, an effect especially strong with the latter treatment. To examine the effect on the number of apical progenitors, we determined by flow cytometry the number of Prominin-expressing (P^+^) cells in the dissociated GE of E18 Gdf15^−/−^ animals after 24 h treatment with GDF15, TBA or SAG, in relation to untreated controls (Fig. 5H). Consistent with our previous observations (Baur et al. 2023), treatment with recombinant GDF15 decreased the pool of P^+^ cells in the mutant GE to about 80% of the cells in untreated samples. In contrast, only treatment with TBA but not with SAG affected P^+^ cells, indicating that differences in cell cycle re-entry but not in SHH signalling cause the increase in apical progenitors in the mutant GE. Taken together these data show that blockade of HDAC6 and ADCY3 recapitulate the effect of exposure of mutant progenitors to GDF15 both with respect to ciliogenesis proliferation and SHH signalling, whereas upregulation of SHH signalling rescues only the defect in cell cycle progression of mutant progenitors.

## Discussion

In this study, we report for the first time that the expression of GDF15 in the V-SVZ contributes to ciliary function of apical neural progenitors and the regulation of their proliferation. Our data show that lack of GDF15 affects the number and the morphology of the organelle in apical progenitors. Previous studies have highlighted that in embryonic apical radial glia, the inheritance of the mother centriole (Wang et al. 2009) or of ciliary remnants (Paridaen, Wilsch-Brauninger, and Huttner 2013) is associated with the rapid generation of primary cilia and the maintenance of apical radial glia character at the expense of the generation of more differentiated progenitors. Alteration in the orientation of the plane of division (Falk et al. 2017) or of ciliogenesis (Hu et al. 2021; Broix et al. 2018) can also affect the number of ciliated progenitors, however, these mutations also affect differentiation and the type of progenitors generated upon cell division. In contrast, despite the increased number of primary cilia in the mutant GE, we did not observe a change in differentiation but only an increase in the number of generated cells (Baur et al. 2023). This indicates that the increased presence of ciliated cells essentially reflects an expansion of the number of ciliated progenitors. Consistent with this hypothesis, we found that not only apically dividing progenitors but also their progeny, including fast proliferating secondary short progenitors and subapical progenitors, were increased in the mutant GE (Baur et al. 2023). Besides the number of cilia, we report here that mutant cilia display a shorter and thicker appearance. Whereas the importance of the latter to the regulation of the cell cycle is unclear, it is well documented that the length of primary cilia affects cell cycle progression in neural progenitors (Li et al. 2011). A negative correlation between cilia and cell cycle is also observed in the adult V-SVZ, where the length of primary cilia is associated with increased permanence into quiescence (Khatri et al. 2014). In the adult V-SVZ cilia are also particularly associated with the apical side of the V-SVZ (Beckervordersandforth et al. 2010; Baur et al. 2022). However, in contrast to the germinal niche of the HP, where primary cilia are essential for the generation and the maintenance of adult NSCs (Han et al. 2008; Breunig et al. 2008; Amador-Arjona et al. 2011), the relevance of primary cilia for the regulation of neurogenesis in the V-SVZ remains elusive, as ablation of primary cilia in this region appears to reduce proliferation and neurogenesis only in the ventral anterior telencephalon (Tong et al. 2014). A possible explanation for these findings is that primary cilia in the adult V-SVZ are mostly present in a subset of apical NSCs (Khatri et al. 2014), which are slowly cycling cells and do not represent the main contributors of adult neurogenesis (Baur et al. 2022).

Besides cilia length, SHH signalling was also affected in mutant progenitors. Previous observations have shown that ablation of primary cilia in the developing telencephalon from mid-development onwards eliminate Gli1 expression around in the periventricular area, which leads to a decrease in the number of Nkx2.1 expressing cells (Tong et al. 2014). However, our data show that lack of GDF15 limited endogenous SHH activation also in the HP, where ADCY3 expression, but not the length of primary cilia was affected, indicating that the effect of GDF15 on SHH is associated. with changes in ADCY3 expression but not the cilia length. Notably, interference with SHH signalling either directly (Komada et al. 2008) or indirectly through modulation of the ciliary function (Wilson et al. 2012) alters cell cycle progression, although the direction of the change, may be affected by temporal cues. Our data here support the conclusion that SHH affects cell cycle progression, as we found that mutant progenitors display impaired endogenous activation of SHH signalling, and that this impairment appears to affect cell cycle progression but not cell cycle re-entry.

We found that the effect of GDF15 on ciliary function is limited to the V-SVZ, thereby providing a likely explanation for the differences of the growth factor in the regulation of proliferation in the two neurogenic regions. Indeed, in two previous studies we have shown that late in development the expression of GDF15 increases in the dentate gyrus and in the VZ of the GE and that in both brain regions GDF15 ablation leads to a decrease of EGFR expression at the cell membrane. However, whereas the changes in EGFR expression are associated with a decrease in the proliferation of hippocampal progenitors, in the GE altered EGFR is not responsible for the increased proliferation of apical progenitors, which leads to a permanent increase of NSCs and ependymal cells in the adult V-SVZ. Increased EGFR expression is a characteristic of proliferating progenitors, including activated NSCs and transit amplifying progenitors in both germinal niches and from late development onwards EGF is the main mitotic for NSCs, making its expression a reliable marker for the identification of these cell populations in both neurogenic regions. However, how EGFR signalling contributes to proliferation is still unclear since the effect of EGFR activation on proliferation both vary according to age and multiple intracellular pathways are activated contributing to different functional outputs including survival and differentiation (Cochard et al. 2021). Indeed, ablation of EGFR or its overactivation have highlighted a role for EGFR signalling in glial differentiation and survival, which is temporally associated with the acquisition of EGFR responsiveness in neural progenitors at late developmental stages (Zhang et al. 2023). Notably, in the rat V-SVZ EGFR has been associated with the organelle and with the microtubules of the mitotic spindle (Danilov et al. 2009), suggesting that changes in ciliogenesis may underlie the differences in expression of the growth factor between WT and mutant cells. However, this is unlikely since although GFRAL and EGFR are expressed in the same progenitors, we could never detect EGFR expression in primary cilia. Most importantly, changes in EGFR expression were also observed in the HP where ciliogenesis was not affected. Notably, it has been shown that HDAC6 affects EGFR trafficking and degradation (Gao, Hubbert, and Yao 2010) and that absence of the deacetylase leads to faster degradation of the receptor. Consistent with this hypothesis, our data indicate HDAC6 activation and/or subcellular localization are different in WT and mutant progenitors, as illustrated by the fact that blockade of HDAC6 affects intracellular acetylation in apical WT but not mutant progenitors and that ciliary length is differentially affected by HDAC6 blockade depending in the two groups of apical progenitors (Sanchez de Diego et al. 2014). However, EGFR expression is also altered in hippocampal progenitors where we could not detect a change in HDAC6, indicating that GDF15 affects the trafficking of the receptor in an additional manner. Notably, *Adcy3* expression was affected also in the mutant HP, which displayed normal cilia. This indicates that local activation of ADCY3 may be necessary to induce the alteration in ciliogenesis. Interestingly we have found that GFRAL is localized to primary cilia only in the GE/V-SVZ, but not in the HP, suggesting the exciting possibility that the receptor may affect the activation of ADCY3 within the ciliary space. However, local clues reflecting intrinsic and extrinsic differences may also be involved. For example, whereas prominin has been used to sort adult NSCs in both neurogenic regions (Walker et al. 2013; Carrillo-Garcia et al. 2010) like GFRAL it is present in primary cilia only in NSCs of the V-SVZ (Walker et al. 2013).

In light of the fact that GDF15 is expressed apically in the lateral ventricle from late development onwards, our data indicate that the growth factor may contribute to the decline in the number of ciliated apical cells and increase in cilia length observed in this region during aging.

## Acknowledgments

K.B. was supported by the Interdisciplinary Center for Neuroscience (IZN) and the Landesgraduiertenförderung (LGF) of the Heidelberg University Graduate Academy. C.C. was supported by contract research from the program “Adult neural stem cell” of the Baden-Württemberg Stiftung.

