## supplementary figures and tables for "Growth/differentiation factor 15 controls primary cilia morphology in the murine ventricular-subventricular zone thereby affecting progenitor proliferation"

### Supplementary Tables

**Supplementary Table S1:** Antibodies used for immunofluorescence. N/A = not available.

| Antigen | Host | Company, Catalog # | Lot # | Concentration |
|---|---|---|---|---|
| ADCY3 | rabbit | Thermo Fisher, PA5-35382 | UL2902981 | 1:500 |
| Arl13b | mouse | UC Davis, 75-287<br>BioLegend, 857602 | 472-1JU-55<br>B323369 | 1:500 |
| GFRAL | sheep | Invitrogen, PA5-47769 | UH2824346A | 1:200 |
| HDAC6 | rabbit | Proteintech, 12834-1-AP | 00053805 | 1:500 |
| Ki67 | rabbit | Abcam, 16667 | GR3313195-18 | 1:100 |
| Tubulin, acetylated | mouse | Sigma Aldrich, T6793 | 017M4806V | 1:1000 |

**Supplementary Table S2:** qPCR probes for TaqMan assays.

| Gene | Assay # |
|---|---|
| Beta-actin | Mm00607939_s1 |
| Adcy3 | Mm00460371_m1 |
| Gli1 | Mm00494654_m1 |
| HDAC6 | Mm00515945_m1 |

### Supplementary Figures

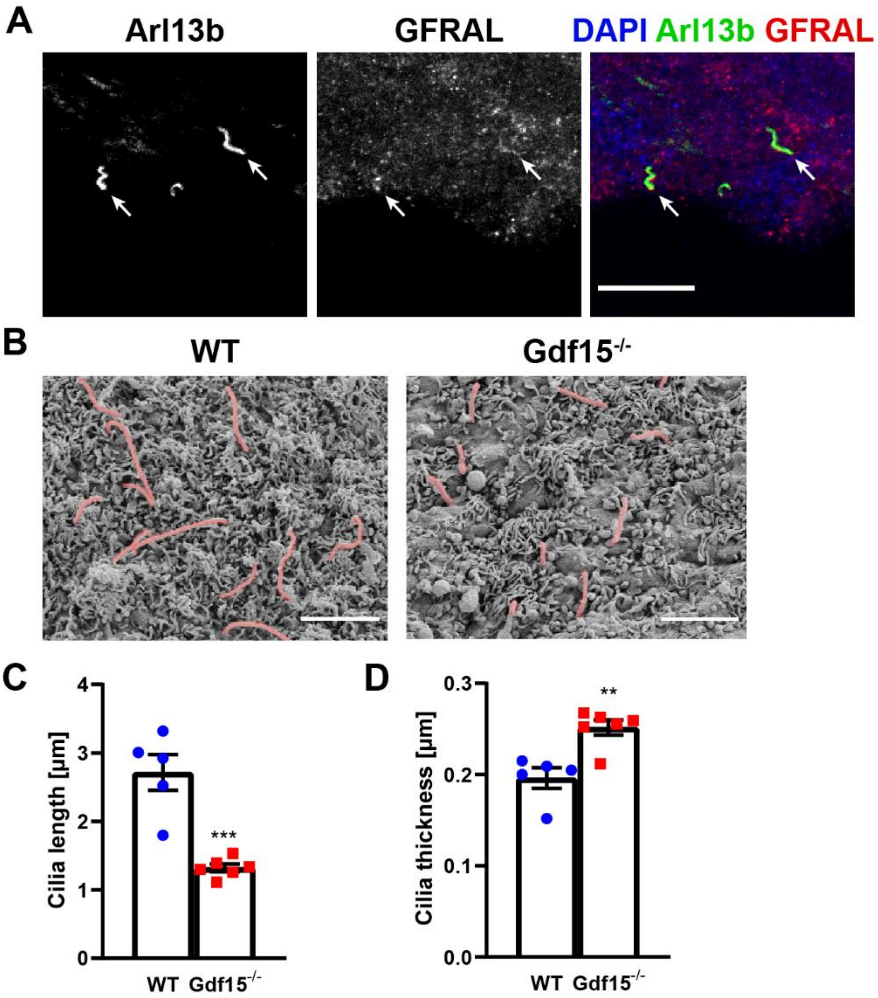


**Supplementary figure S1: Scanning electron microscopy of E18 GE primary cilia.**

(A) Immunofluorescent micrographs of coronal sections of adult WT mice, labelled for cilia marker Arl13b (green) and GFRAL (red). DAPI was used for nuclear counterstain. Scale bar = 20 µm.

(B) Representative images of primary cilia (red) at the apical side of the E18 ganglionic eminence of the given genotype upon scanning electron microscopy (ScEM). Scale bars = 2 µm.

(C, D) Quantitative analysis of cilia length (C) and thickness (D) in ScEM samples. Bars indicate mean ± SEM. Each dot represents individual animals. * indicates significance from WT (**$p<0.01$; ***$p<0.001$).

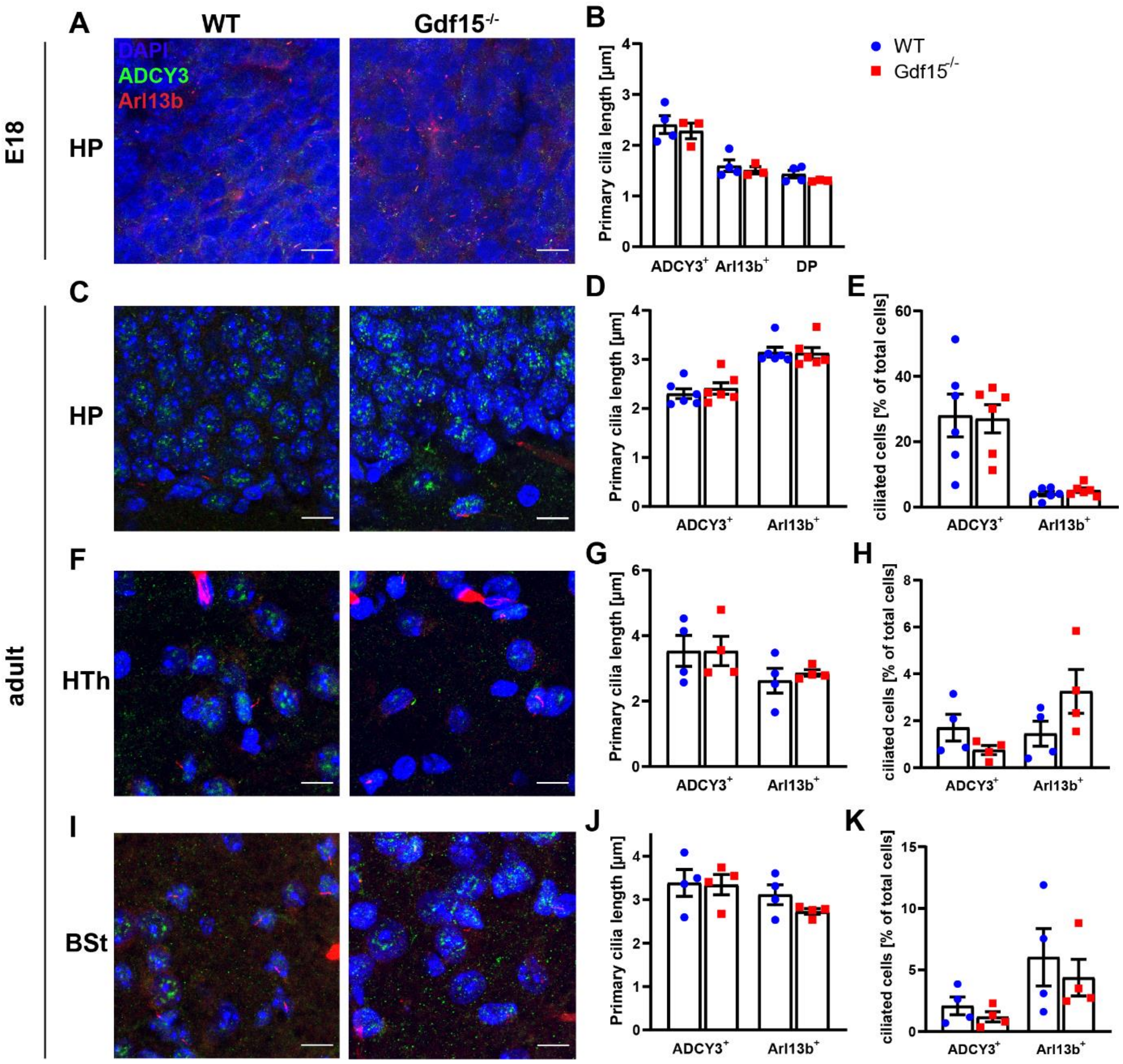


**Supplementary figure S2: GDF15 does not affect cilia morphology and number in other brain regions**.

(A, C, F, I) Representative confocal micrographs illustrating primary cilia in the dentate gyrus of the hippocampus (HP; A, C) and hypothalamus (Hth; F) and brain stem (Bst; I) obtained from coronal sections of E18 (A) and adult (C, F, I) brains upon double immunostaining for ciliary marker Arl13b (blue) and ADCY3 (green), with DAPI (blue) for nuclear counterstaining. Scale bars = 10 µm.

(B, D, E, G, H, J, K) Graphs illustrate quantification of primary cilia length (B, D, G, J) and thickness (E, H, K) in each region according to genotype and cilia marker. Bars indicate mean ± SEM. Each data point represents summarized data from one individual animal.

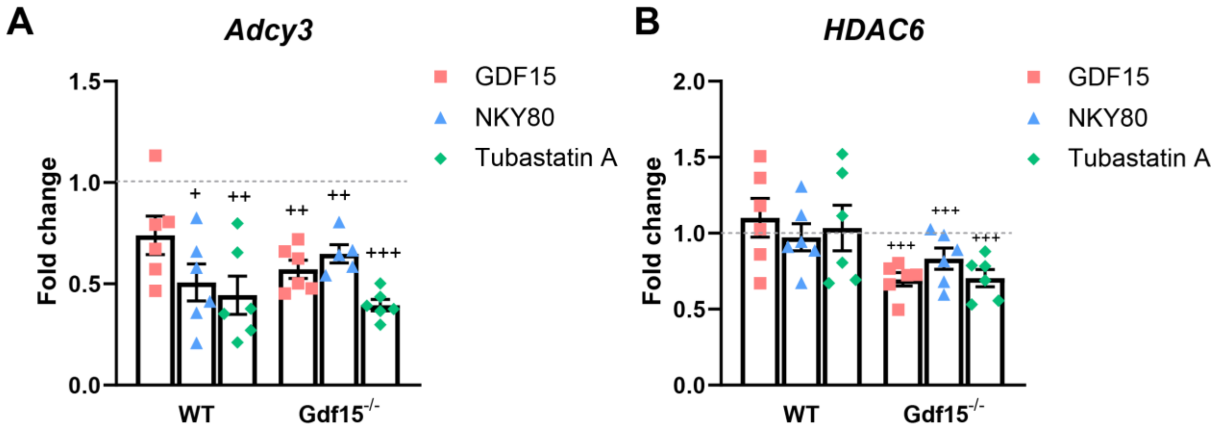


**Supplementary figure S3: Effect of inhibition on *Adcy3* and *Hdac6* mRNA levels.**

Fold change in the expression of *Adcy3* (A) and *Hdac6* (B) mRNA in the whole E18 GE according to the genotype and treatment. Values are normalized to respective untreated controls (dashed lines). Bars indicate mean ± SEM. Each data point represents summarized data from one individual animal. $^{+}$ indicates significance from control. ($^{+}p<0.05$, $^{++}p<0.01$, $^{+++}p<0.001$).
